# An immunoinformatics approach to study the epitopes contributed by Nsp13 of SARS-CoV-2

**DOI:** 10.1101/2021.04.02.438155

**Authors:** Sushant Kumar, Gajendra Kumar Azad

## Abstract

The on-going coronavirus disease-19 (COVID-19) pandemic caused by SARS-CoV-2 has infected hundreds of millions of people and killed more than two million people worldwide. Currently, there are no effective drugs available for treating SARS-CoV-2 infections; however, vaccines are now being administered worldwide to control this virus. In this study, we have studied SARS-CoV-2 helicase, Nsp13, which is critical for viral replication. We compared the Nsp13 sequences reported from India with the first reported sequence from Wuhan province, China to identify and characterize the mutations occurring in this protein. To correlate the functional impact of these mutations, we characterised the most prominent B cell and T cell epitopes contributed by Nsp13. Our data revealed twenty-one epitopes, which exhibited high antigenicity, stability and interactions with MHC class-I and class-II molecules. Subsequently, the physiochemical properties of these epitopes were also analysed. Furthermore, several of these Nsp13 epitopes harbour mutations, which were further characterised by secondary structure and per-residue disorderness, stability and dynamicity predictions. Altogether, we report the candidate epitopes of Nsp13 that may help the scientific community to understand the evolution of SARS-CoV-2 variants and their probable implications.

## INTRODUCTION

The coronavirus disease 2019 (COVID-19) disease was first reported from Wuhan province, China in the late 2019 [1–3]. The causative agent of this disease was identified as Severe Acute Respiratory Syndrome-Corona Virus-2 (SARS-CoV-2), which shares considerable similarity with previously known coronavirus, SARS-CoV [4]. Coronaviruses are a group of RNA viruses, which can infect diverse animals, including humans [5]. Earlier reports have shown that at least six different coronaviruses are known to infect humans, including CoV-229E, CoV-OC43, CoV-NL63, CoV-HKU1, SARS-CoV, and MERS-CoV [6]. Four of them do not cause any serious health implication on humans including CoV-229E, CoV-OC43, CoV-NL63, and CoV-HKU1 but serious respiratory issues have been linked to SARS-CoV and MERS-CoV infection [7]. The SARS-CoV-2 is the seventh coronavirus to infect humans and also causes mild to severe respiratory illness in the infected individuals and reported to cause severe symptoms in immune-compromised patients. The SARS-CoV-2 rapidly spread worldwide within few months and become one of the worst pandemic ever reported in human history [8]. Still this virus is spreading and already triggered second and third wave of infections in many countries. As of 18^th^ March 2021, more than 120 million cases of COVID-19 were reported worldwide with more than 2.6 million deaths.

SARS-CoV-2 genome is comprised of a positive sense single stranded RNA of approximately 29kb [9]. Its genome encodes four structural, sixteen non-structural and nine accessory proteins [10]. The non-structural proteins are involved in the maintenance of functional integrity of the virus and also required for infection and virus particle formation. Numerous RNA viruses have been found to encode their own RNA helicases, which are usually indispensable components of the RNA replication complexes [11,12] and recognized as an ideal targets for developing antivirals [13]. Nsp13, an RNA helicase, plays an important role in the folding of its RNA elements or unwinding of double stranded RNAs. Previous studies have shown that SARS-CoV Nsp13 has an NTPase and helicase activity belonging to helicase superfamily-1 [14]. The SARS-CoV-2 Nsp13 helicase shares a 99.8% sequence identity to SARS-CoV (SARS) Nsp13 helicase [15]. Nsp13 helicase is a critical component for viral replication and shares the highest sequence conservation across the CoV family, highlighting their importance for viral viability. RNA dependent RNA polymerase (RdRp) and Nsp13 are required for viral replication, therefore, both of these vital enzyme represents a promising target for anti-CoV drug development [16].

In order to understand the host response towards SARS-CoV-2, we performed in-silico study to characterize Nsp13. We used several bioinformatic tools to predict potential epitopes of Nsp13 and characterised them by studying several immunological parameters. Further, we systematically characterized the mutations in Nsp13 reported from India and their impacts on epitopes are discussed.

## MATERIALS AND METHODS

### Nsp13 sequence retrieval and multiple sequence alignment (MSA)

The Orf1ab of SARS-CoV-2 contains Nsp1-16 as a single polypeptide which is proteolytically cleaved into individual proteins. The Orf1ab sequences used in this study were downloaded from NCBI-virus database. Till 3^rd^ March 2021, 651 sequences of Orf1ab have been reported from India. We have downloaded all these sequences and their protein accession number are listed in the supplementary table 1. The Nsp13 sequences were retrieved from their respective Orf1ab and used for analysis. The Orf1ab sequence reported from Wuhan, China was used as a reference sequence (Protein accession number: YP_009724389) for MSA studies using Clustal Omega tool [17] as described earlier [18].

### B cell epitope prediction

Linear B cell epitopes were predicted using IEDB (Immune Epitope Database and Analysis Resource) [19], an online server tool based on Bepipred linear epitope prediction method (at the threshold value of 0.350). IEDB prediction tool was also used to predict the Nsp13 immunological parameters including the antigenicity, accessibility, flexibility, hydropathicity and beta turn. These standard parameters were estimated by Chou and Fasman beta-turn prediction algorithm, Emini surface accessibility server tool, Karplus and Schulz flexibility prediction tool, Kolaskar and Tongaonkar antigenicity and Parker hydrophilicity prediction algorithms, respectively [20,21]. The DiscoTope 2.0 was used for prediction of Discontinuous B cell epitope using threshold value set at −5.5 and its three dimensional structure were represented by AutoDock software. Antigenicity and allergenicity of B cell epitopes were predicted by Vaxijen 2.0 [22] and AllergenFP v.1.0 [23] servers, respectively.

### T cell epitope prediction

On the surface of antigenic presenting cell, T cell epitopes are presented where they are attached to Major Histocompatibility Complex (MHC) molecule. MHC class I and class II molecules were predicted as follows-

MHC class I molecule: IEDB webserver based on NetMHCpanEL 4.1 was used for prediction of MHC class I molecules [24]. For this prediction, we selected HLA reference alleles (a total of 54 alleles) having epitope length of 9 or 10 mers. We finally sorted 9 mers conserved epitopes that show maximum binding interaction at IC50 < 200nm. Antigenicity of all selected epitopes was predicted from Vaxijen 2.0 webserver, whereas allergenicity was predicted by AllergenFP v.1.0.

MHC class II molecules: IEDB recommended 2.22 prediction method was used to predict MHC class II epitopes. For this prediction, we selected the seven standard reference alleles having maximum length of 11 mers. Subsequently, we sorted most conserved 9 mers epitopes that exhibited maximum binding interaction with other alleles. Antigenicity of all selected epitopes was predicted from Vaxijen 2.0 webserver, whereas allergenicity was predicted by AllergenFP v.1.0.

### Physiological profiling of T cell epitopes-

The characterisation of selected MHC class I and MHC class II epitopes were performed by several webservers. The physiological parameters including toxicity, hydrophobicity, hydropathicity, charge PI and molecular weight were calculated by Toxinpred tool [25]. Another webserver tool HLP [26] was used to predict half-life, surface accessibility, flexibility and polarity of the selected epitopes.

### Structure modelling, secondary structure analysis and protein disorder prediction

The locations of identified mutations were highlighted in the three-dimensional structure of Nsp13 using UCSF Chimera program [27]. The Nsp13 RCSB-ID: 6ZSL was used for the structural representation. The prediction of secondary structure was performed by CFSSP webserver [28] as described earlier [29]. The input sequence was uploaded on this server, which provides the secondary structure in terms of alpha-helix, beta-sheet and turns. The prediction of per residue intrinsic disorder predisposition contributed by each residue of input Nsp13 polypeptide sequence was performed by PONDR-VSL2 webserver [30]. The value above 0.5 is considered ordered while the value below 0.5 is considered disordered.

### Stability of protein structure

The stability of protein structure was predicted by DynaMut webserver [31]. The reported structure of Nsp13 (RCSB-ID: 6ZSL) was used to predict the impact if mutation on the stability of protein. First, the protein structure was uploaded onto the webserver, followed by providing the details of the mutation for analysis as described earlier [32]. This webserver predict the differences in free energy between the wild type and mutant protein.

## RESULTS

### Identification of variations in nsp13 protein among Indian isolates

We compared the Nsp13 sequences reported from India with the first reported sequence from Wuhan province, China using Clustal omega tool. Our analysis revealed twenty-seven mutations have taken place among Indian isolates, which is distributed all over the Nsp13 protein as shown in figure 1A. Next, we observed the location of these mutations in the three-dimensional structure of Nsp13 using UCSC Chimera tool as shown in figure 1B. We also looked at the polarity change and charge alteration due to the mutation in Nsp13, our data revealed that most of mutation does not led to any change (neutral to neutral) however, at two positions it changed from acidic to neutral (E142V and E168A) while at three locations it altered from basic to neutral (H164Y, R392C and R442Q) as shown in table 1.

**Figure 1:**
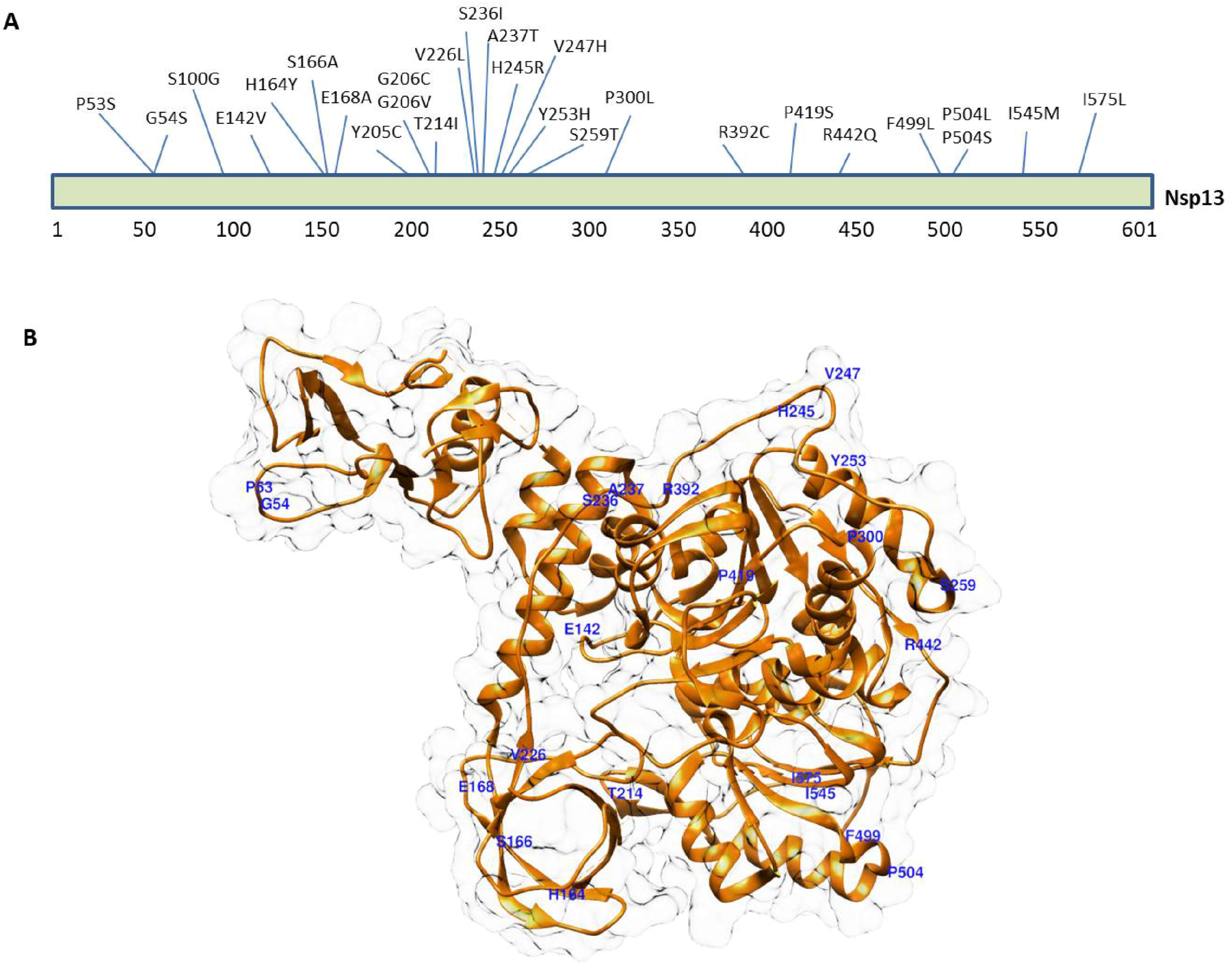
The diagrammatic representation of Nsp13 of SARS-CoV-2 demonstrating the details and location of mutations identified in this study. The SARS-CoV-2 Nsp13 protein is comprised of 601 amino acid residues. B) Structural representation of three dimensional structure of the Nsp13 (RCSB protein ID-6ZSL) showing the location of mutations (single amino acid letter code) along with its respective position in polypeptide sequence. The structural data was obtained from UCSF chimera tool.

**Table 1:**
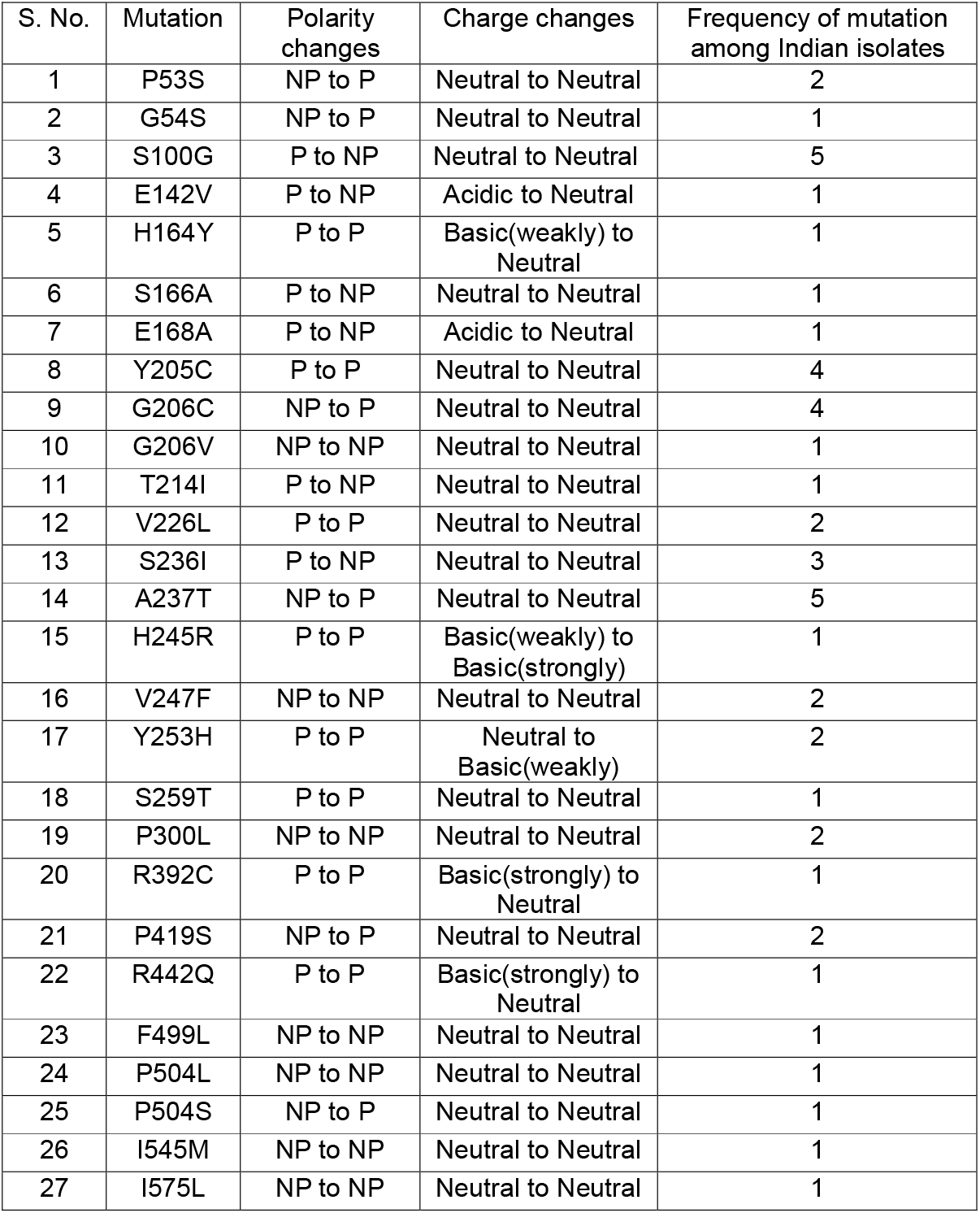
The table show the location and details of mutations identified in Nsp13 of SARS-CoV-2. The frequency of each mutation along with polarity and charge changes are also indicated.

### Prediction of B cell epitope contributed by Nsp13

IEDB webserver tool was used for predicting the continuous B-cell epitopes of Nsp13 (figure 2A). Our analysis revealed seven best antigenic epitopes that were more than 8 amino acid residue in length (figure 2B). Subsequently, these epitopes were further characterised by analysing various parameters, including vaxijen score, allergenicity, and toxicity (figure 2B). Our prediction data revealed that all of them are non-toxic however, 5 peptides (CNAPGCDVT, CVGSDNVT, VGKPRPPLN, TFEKGDYGDA and GDPAQLPAP) possess alleregenicity properties and two peptides are non-allergen (TQTVDSSQGSEY and STLQGPPGTGKS). Similarly, three of them exhibited antigenic property while four are non-antigenic (figure 2B).

**Figure 2:**
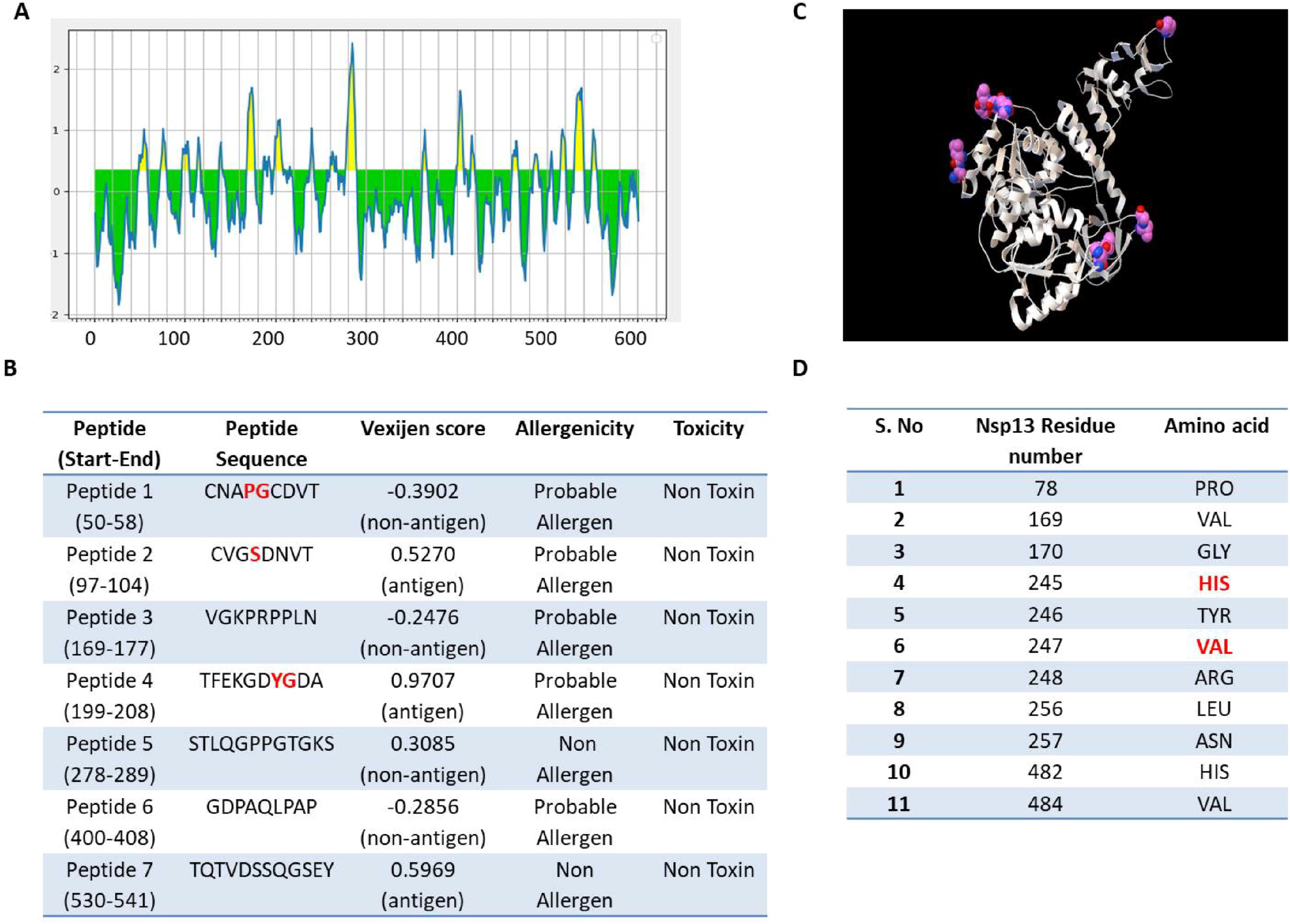
Prediction of B-cell epitopes of Nsp13. Linear continuous epitopes, A) The Y-axis of the graph corresponds to BepiPred score, while the X-axis depicts the Nsp13 residue positions in the sequence. The yellow area of the graph corresponds to the part of the protein having higher probability to be part of the epitope. B) The top seven peptides of Nsp13 having at least 8 amino acids are shown with vaxijen score. The red font shows the location of mutant residues in the epitope sequence. C) Prediction of discontinuous B-cell epitopes. The position of each predicted epitope on the 3D structural surface of Nsp13 was denoted using Autodock. D) The location and identity of each discontinuous epitopes of Nsp13.

Subsequently, we predicted the B cell epitopes of Nsp13 based on its three dimensional structure using DiscoTop 2.0 webserver tool [33]. Our analysis revealed the eleven discontinuous epitopes of Nsp13 having high score. The location of these epitopes are highlighted on the 3D structure of Nsp13 (figure 2C) and their additional details are shown in figure 2D. Altogether, our data revealed B –cell epitopes contributed by Nsp13.

### Analysis of essential parameters of B cell epitopes

Subsequently, all essential parameters of B Cell epitopes including Beta turn, accessibility of surface, flexibility, antigenicity and hydrophilicity were also calculated for Nsp13 (figure 3). Chou and Fasman’s beta-turn prediction algorithm (with threshold 0.984) resulted in a minimum score of 0.677 and a maximum score of 1.384, and our selected peptide showed a propensity score of 1.164 (figure 3A). Emini surface accessibility prediction tool (with threshold 1.000) resulted in a minimum score of 0.036 and a maximum score of 4.888, and our selected peptide scored 1.33 in surface accessibility (figure 3B). Karplus and Schulz’s flexibility prediction tool (with threshold 0.989) resulted in a minimum score of 0.904 and a maximum score of 1.150, and our selected peptide showed 1.033 in flexibility score (figure 3C). Kolaskar and Tongaonkar antigenicity scale (with threshold 1.052) resulted in a minimum score of 0.893 and a maximum score of 1.284, and our selected peptide showed an antigenicity score of 1.0163 (figure 3D). The parker hydrophilicity prediction algorithms (with threshold 1.325) resulted in a minimum score of −3.714 and a maximum score of 7.000, and our selected peptide showed 1.33 in hydrophilicity score (figure 3E).

**Figure 3:**
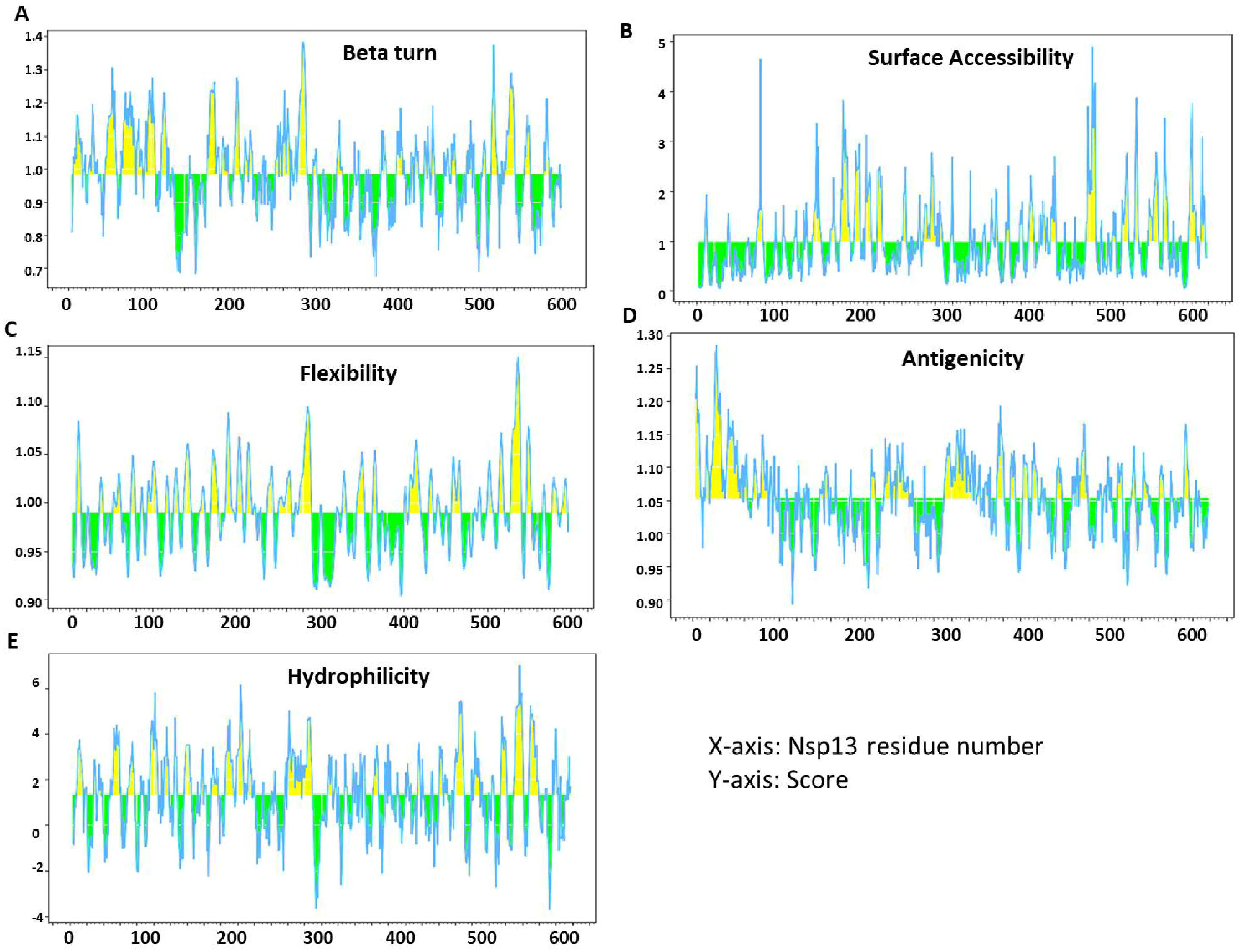
Recognition of B cell epitopes. A) Antigenic determinants of Nsp13 were predicted using Kolaskar and Tongaonkar, B) Hydrophilicity of Nsp13 was predicted by Parker hydrophilicity, C) Surface accessibility analysis shown on Emini surface accessibility scale, D) β variants of structural polyproteins as predicted by Chou and Fasman β metamorphosis prediction, E) Flexibility analysis on Karplus and Schultz flexibility scale.

### Prediction of T cell epitopes contributed by Nsp13 protein

The IEDB webserver tool was used for prediction of Cytotoxic T-Lymphocyte epitopes and its interaction with MHC class I molecules. Our analysis with Nsp13 revealed 8 potential T cell epitopes (table 2). NetMHCpanEL 4.1 MHC-class I binding prediction tools was used to predict the binding of these epitopes with MHC class I molecules with high affinity are listed in table 2. The peptides FAIGLALYY from start (11) to end (19) had highest immunogenicity and affinity to interact with 9 alleles (HLA-B*35:01, HLA-A*26:01, HLA-B*53:01, HLA-A*01:01, HLA-A*30:02, HLA-B*58:01, HLA-B*15:01, HLA-A*68:01, HLA-B*57:01) and also showed allergenicity (table 2). The allergenicity of these epitopes was predicted by Allergen FP tool, which categorises peptides into allergen/non-allergen based on the Tanimoto coefficients.

**Table 2:**
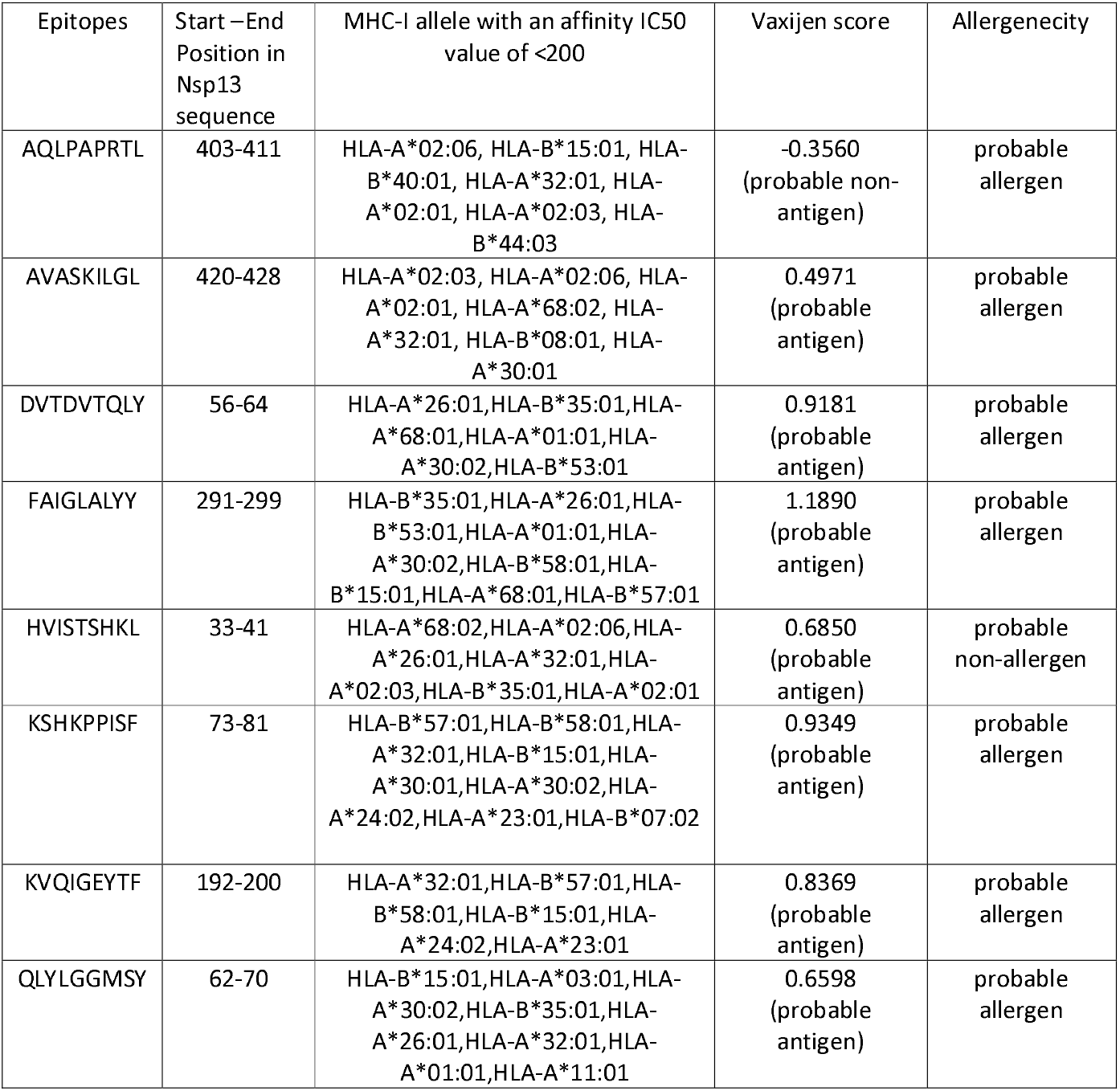
The table show the location and details of MHC-Class-I peptides of Nsp13. The MHC-I interaction with top most allele (affinity IC50 value of <200) are mentioned in the table along with Vaxijen score and allergenicity.

Similarly, the prediction of Helper T-Lymphocyte epitopes of Nsp13 and its interaction with MHC Class II molecules was predicted by IEDB webserver (based on IEDB recommended 2.22 method). We selected top 6 epitopes that exhibited maximum binding affinity with MHC class II molecules (table 3). Our analysis revealed that only one epitope (HKLVLSVNP) possesses both antigenicity and allergenicity properties and affinity to interact with 5 alleles as shown in table 3 (HLA-DRB1 *15:01, HLA-DRB1 *03:01, HLA-DRB1 *07:01, HLA-DRB3*01:01, HLA-DRB5*01:01). The epitopes EHYVRITGL had highest immunogenicity and affinity to interact with 4 alleles (HLA-DRB5*01:01, HLA-DRB1*15:01, HLA-DRB4*01:01, HLA-DRB1*07:01) (table 3). The allergenicity of these epitopes were also predicted based on Tanimoto coefficients.

**Table 3:**
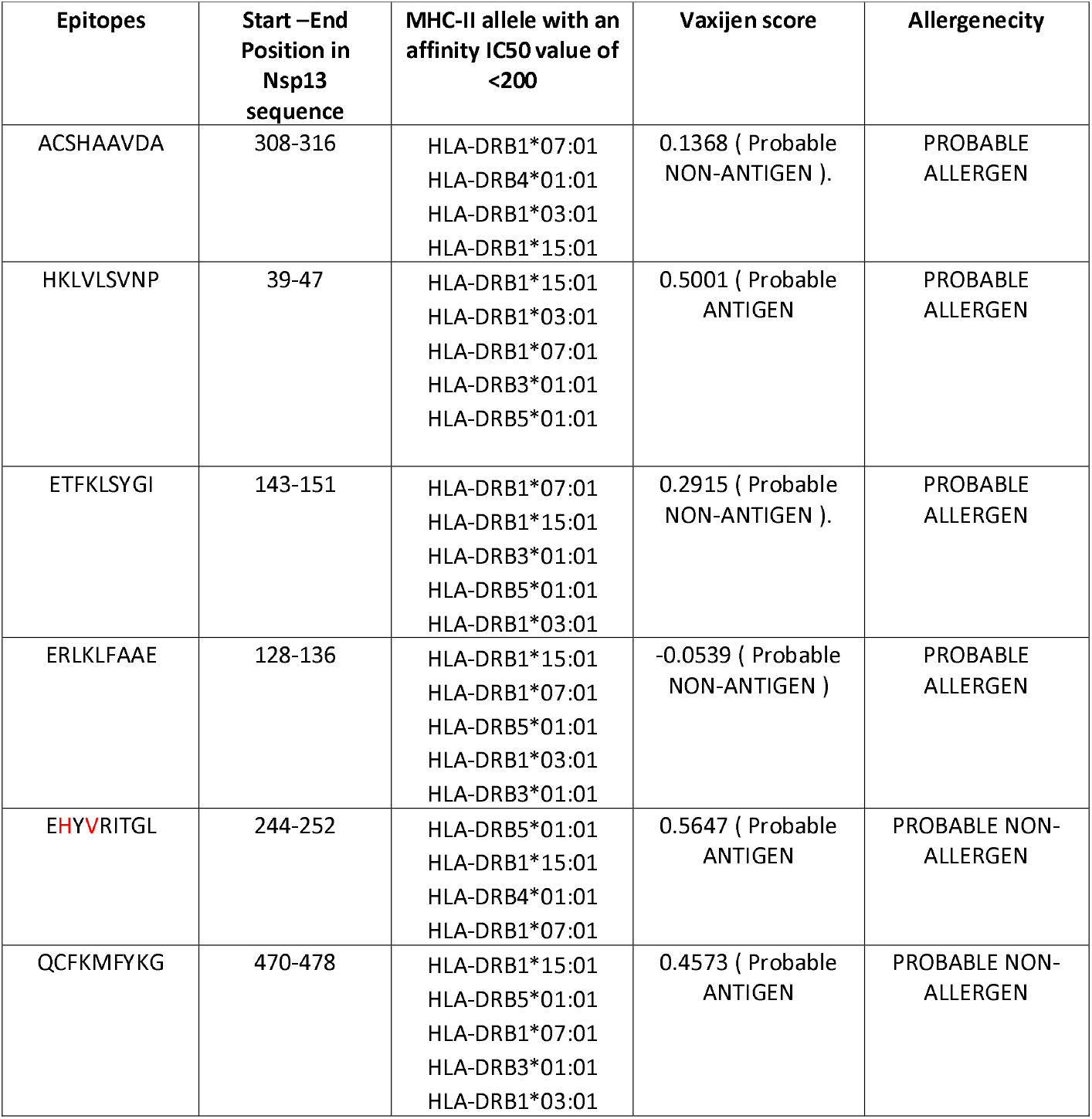
The table show the location and details of MHC-Class-II peptides of Nsp13. The MHC-II interaction with top most allele (affinity IC50 value of <200) are mentioned in the table along with Vaxijen score and allergenicity.

### Physiochemical profiling of promising T-cell epitopes

To substantiate our data, we examined several vital physiochemical features of the promising T cell epitopes. The half-life of MHC class I and II epitopes were calculated and our data revealed that the maximum half-life was observed for HVISTSHKL (MHC class I) and ETFKLSYGI exhibited for MHC class II molecule (table 4). The toxicity prediction was performed by ToxinPred tool show that all analysed molecules were non-toxic (table 4). We further analysed hydrophobicity, hydropathicity, hydrophilicity, polarity, charge, flexibility, pI, molecular weight and surface accessibility of both MHC class I and II molecules (table 4).

**Table 4:**
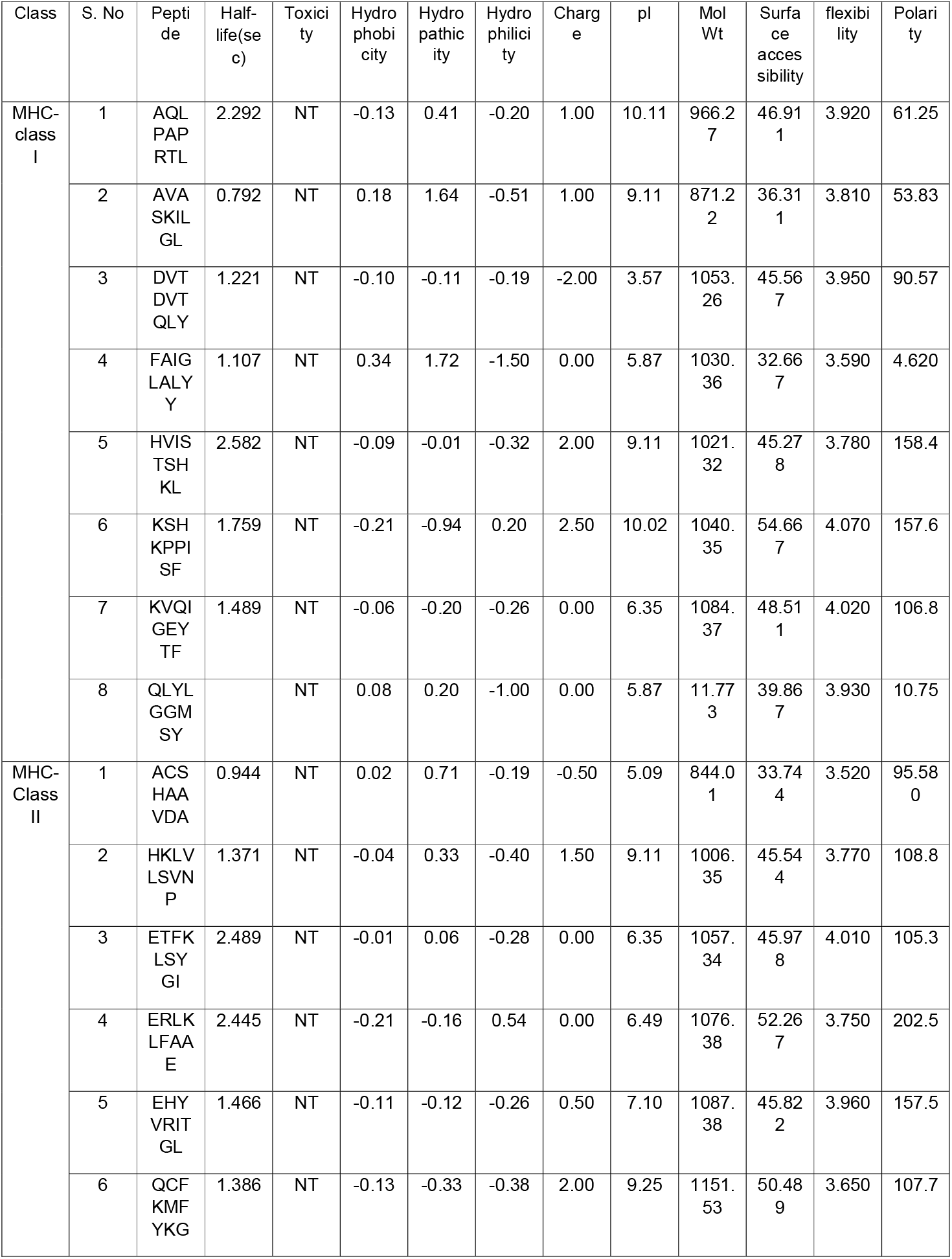
The table show the physiochemical properties of MHC-I and MHC-II peptides of Nsp13.

### The secondary structure prediction revealed several mutations that might lead to structural change of Nsp13

To understand the effect of mutation on Nsp13 protein, we performed secondary structure prediction by CFFSP webserver. This prediction tool identifies the variation in secondary structure contributed by the mutant residue. Our data revealed that out of 27 mutations, only 12 show alterations in secondary structure (figure 4). The minor variations were observed at positions E142V, H164Y, G206V, T214I, S259T, R392C, P419S, and R499L (figure 4A-D, G, I, J and K). S236I and A237T mutation lead to replacement of helical structure into beta-sheet (figure 4E and F). The coiled coil and turn structure were shifted to helix and beta-sheet in P300L and P504L mutants (figure 4H and L).

**Figure 4:**
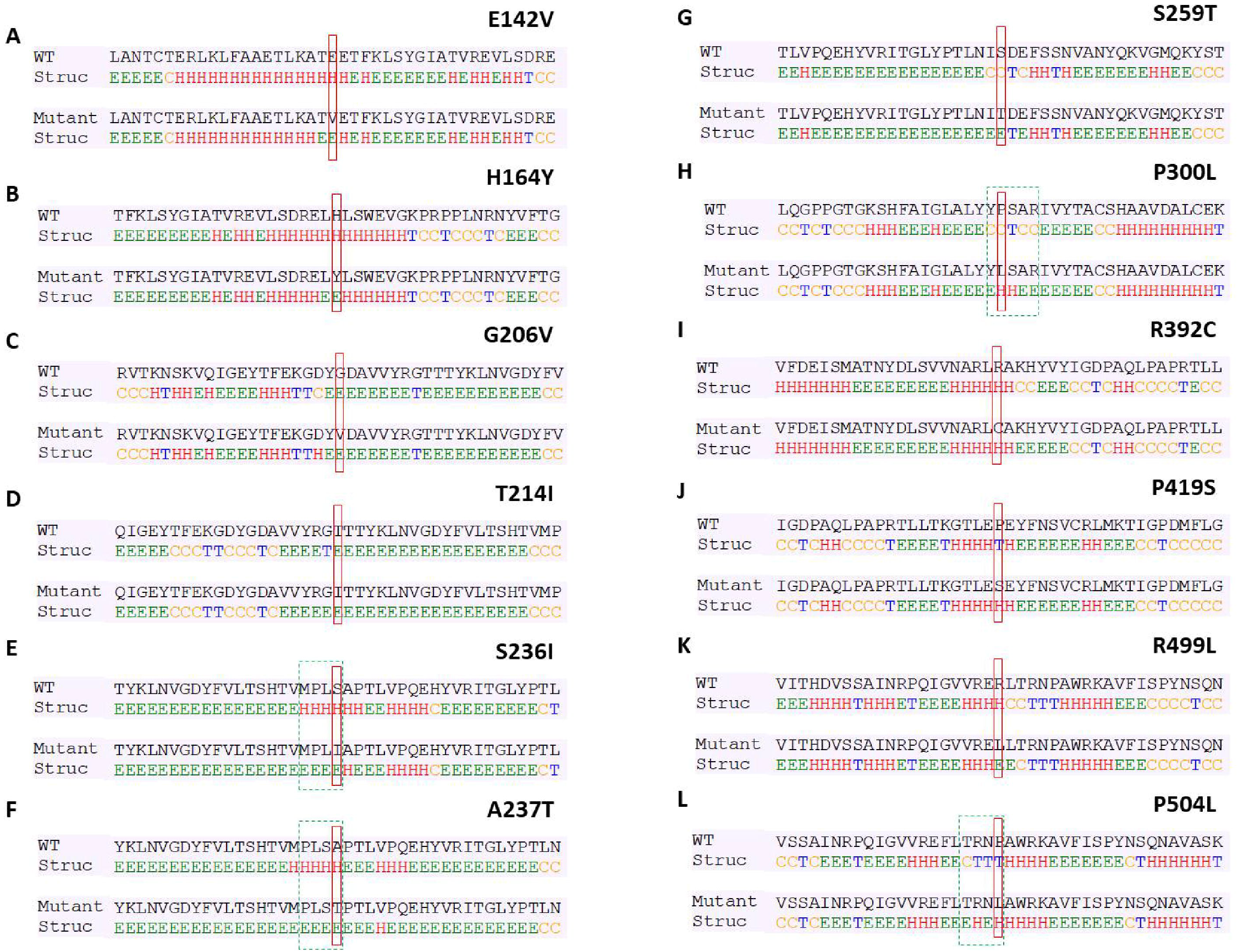
Effect of identified mutations of Nsp13 on its secondary structure. Panel (i) represents the wild type sequence (Wuhan isolate) and panel (ii) represents mutant sequence (Indian isolate). (A-L) each panel represents the individual mutant as depicted. In each panel the contribution of individual amino acid to the secondary structure of Nsp13 protein are shown (H-helix, C-coiled-Coil, T-turn, E-beta sheet). The data was generated from CFSSP tool. The red box in each panel highlights the location of the wild-type and mutant residue.

### Protein disorder predictions due to Mutations in Nsp13 protein

Disordered regions (DRs) are defined as entire proteins or regions of proteins that lack a fixed tertiary structure. Alteration of amino acid sequences in polypeptide chain due to the mutation might causes change in protein disorder. Here, we used PONDR-VSL2 webserver to measure the protein disorder of those mutants that showed alteration in secondary structure. Our analysis revealed that ten mutations E142V, H164Y, S166A, E168A, G206V, G206C, T214I, S236I, R392C and P504L (figure-5A, B, C, D, E, F, G, H, J and K) decreased the protein disorder, while two mutations Y253H and I545M (figure-5I and L) led to increase in protein disorder. Altogether, our data revealed alteration in protein disorder due to the mutation in Nsp13 of SARS-CoV-2.

**Figure 5:**
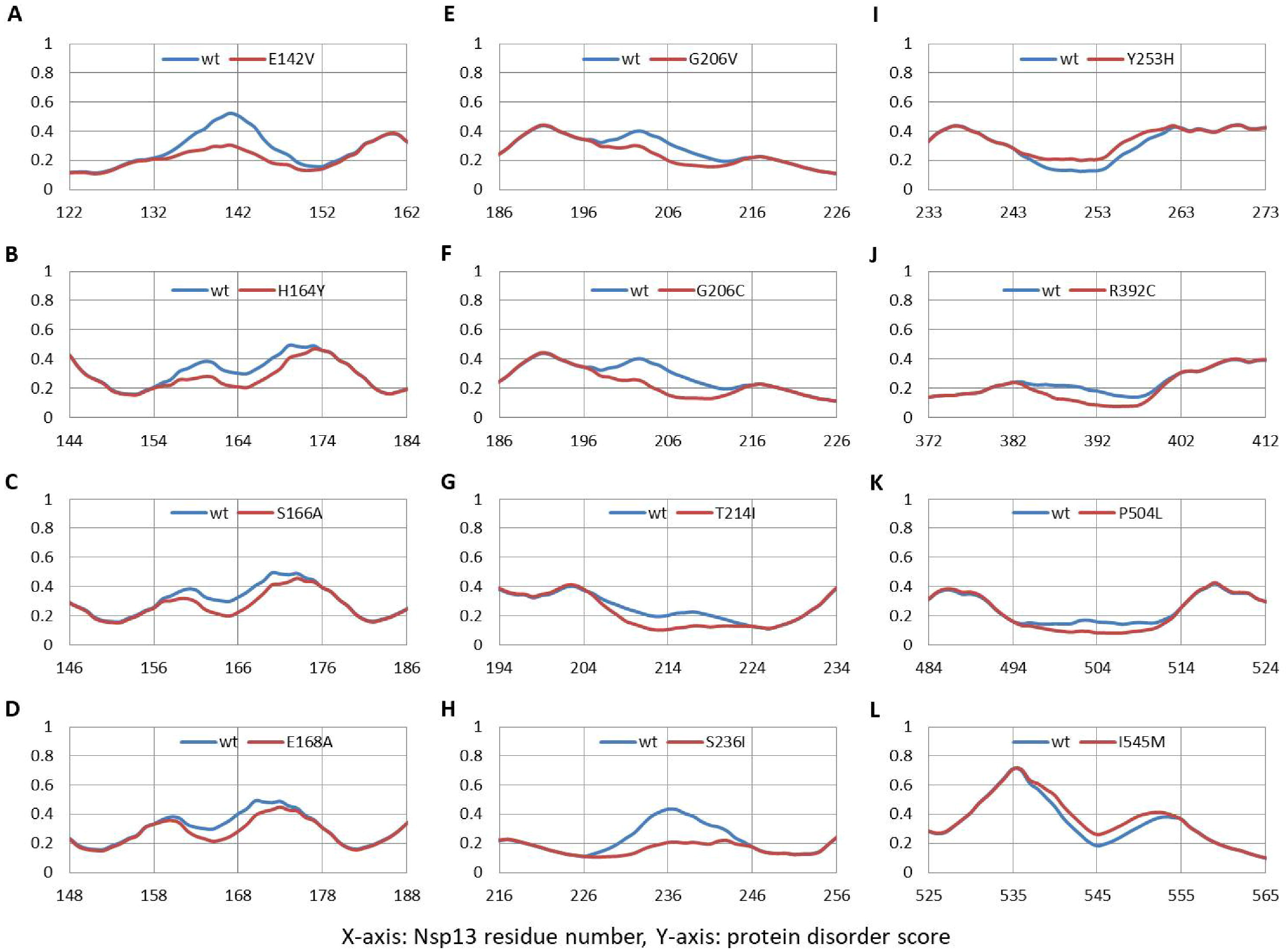
The comparisons of the per residue intrinsic disorder predisposition of Nsp13. A disorder threshold is indicated at score = 0.5, residues/regions with the disorder scores >0.5 are considered as disordered. (A-L) Each panel comparatively represents the disorder parameters of wild type and mutant Nsp13 polypeptide sequence.

### Mutations cause alteration in dynamic stability of Nsp13

We used DynaMut program to predict the effect of mutations on the stability of the protein and calculated the differences in free energy (ΔΔG) between wild type and mutants. The positive ΔΔG corresponds to increase in stability while negative ΔΔG corresponds to decrease in stability. Our analysed data revealed the noticeable increase or decrease in free energy in various mutations as shown in table 5. The maximum positive ΔΔG was observed for H164Y (0.974 kcal/mol) and the maximum negative ΔΔG was obtained for G54S (−1.694kcal/mol) (table 5). Similarly, we also measured the change in vibrational entropy energy (ΔΔSVibENCoM) between the wild type and the mutants. The highest positive ΔΔSVibENCoM was Obtained for P53S (0.504kcal.mol-1.K-1) and negative ΔΔSVibENCoM was obtained for P419S (−0.661 kcal.mol-1.K-1) (table 5). Altogether, the data obtained from ΔΔG and ΔΔSVibENCoM suggests that the mutations identified in this study can influence Nsp13 protein stability and dynamicity.

**Table 5:**
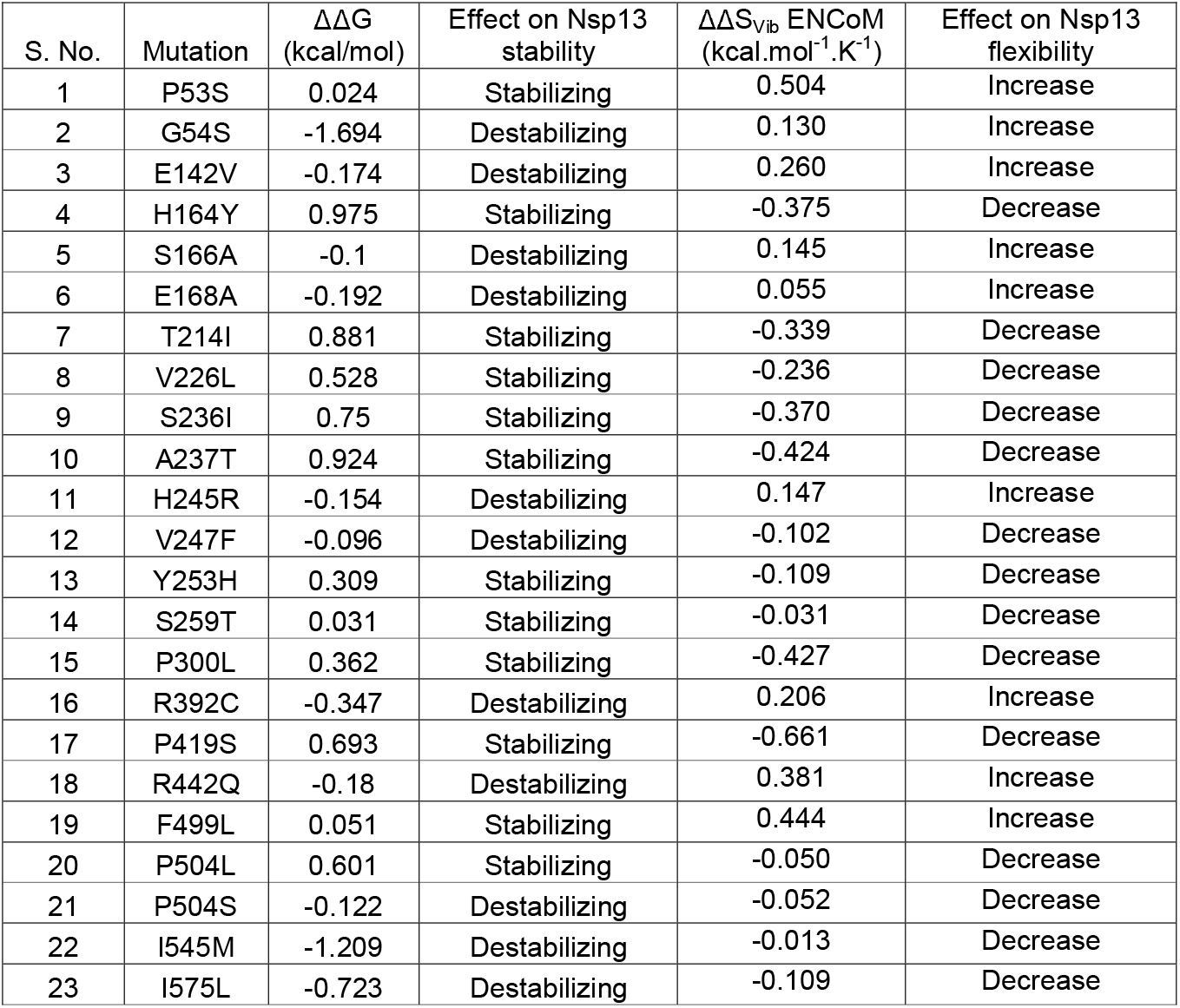
The table show the ΔΔG and ΔΔSvib ENCoM of the mutants present in Nsp13. DynaMut program was used to calculate both parameters.

## DISCUSSIONS

In this study, we examined the physiochemical characteristics of the Nsp13, a non-structural protein of SARS-CoV-2. We studied the high rank B cell and T cell epitope candidates based on the immune-informatics tools to identify Nsp13 epitopes that could regulate host immune responses. The immunoinformatics approaches have been used to study the epitopes from several viruses and those information’s were used to understand the immune response of viruses [34,35]. We used several physiological parameters including structural protrusion, antigenicity, flexibility, surface accessibility, hydrophilicity of Nsp13 were assessed to predict potential B-cell epitopes. IEDB resource tool was used to predict B-cell linear epitopes of at least eight amino acid residues. Our analysis revealed that under these conditions, seven linear B-cell epitopes were predicted that are at least 8 amino acids in length and are of non-toxic. The discontinuous epitopes comprised of the residues that might be separated in linear sequence, however, in 3D structure they are in close proximity [36]. Subsequently, the candidate epitopes were further characterised by the tools that can predict various physicochemical properties. Our data revealed that among B-cell epitopes the ‘TFEKGDYGDA’ peptide exhibited a strong stable immunogenic property as shown by its highest vaxijen score (.9707) and non-toxic. Furthermore, eight MHC class-I and six MHC class-II binding T-cell epitopes assessed as highly antigenic and also predicted to interact with several HLA alleles. Detailed analysis revealed that the best vaxijen score of 1.18 was found for ‘FAIGLALYY’ peptide for MHC class-I moelcule, and for MHC class-II molecules, ‘HKLVLSVNP’ has a maximum vaxijen score of 0.50 epitopes.

It has been well established that the replication of coronaviruses are error-prone that lead to creation of highly diverse genotype variants. Our study revealed a considerable alteration in stability and dynamicity due to mutations at various positions of Nsp13 that might alter its function. In-silico analyses were performed to identify and characterise the mutations occurring in Nsp13. Our data demonstrate that seven of the identified mutations reside in those epitopes that include P53, G54, S100, Y205, G206, H245 and V247 positions (figure 2), which can help the SARS-CoV-2 variants to elicit distinct immune response compared to the wild-type SARS-CoV-2. Among these G206V also led to alteration in secondary structure (figure 4) and protein disorder parameters (figure 5). This proposed consequence due to variations in Nsp13 epitopes are substantiated by several findings with similar observations [37–39]. In short, these results could provide some insights in the understanding of Nsp13 epitopes, which is essential for evaluating the immunogenicity and virulence of SARS-CoV-2.

## ACKNOWLDEGEMENTS

We would like to acknowledge the Department of Zoology, Patna University, Patna, Bihar (India) for providing infrastructural support for this study.

---

Supplementary table 1: List of protein accession number used in this study

